# Multi-condition and multi-modal temporal profile inference during mouse embryonic development

**DOI:** 10.1101/2024.03.03.583179

**Authors:** Ran Zhang, Chengxiang Qiu, Gala Filippova, Gang Li, Jay Shendure, Jean-Philippe Vert, Xinxian Deng, Christine Disteche, William Stafford Noble

**Affiliations:** Department of Genome Sciences, University of Washington; eScience Institute, University of Washington; Brotman Baty Institute for Precision Medicine, University of Washington; Howard Hughes Medical Institute; Allen Center for Cell Lineage Tracing; Owkin; Department of Pathology, University of Washington; Paul G. Allen School of Computer Science and Engineering, University of Washington

## Abstract

The emergence of single-cell time-series datasets enables modeling of changes in various types of cellular profiles over time. However, due to the disruptive nature of single-cell measurements, it is impossible to capture the full temporal trajectory of a particular cell. Furthermore, single-cell profiles can be collected at mismatched time points across different conditions (e.g., sex, batch, disease) and data modalities (e.g., scRNA-seq, scATAC-seq), which makes modeling challenging. Here we propose a joint modeling framework, Sunbear, for integrating multi-condition and multi-modal single-cell profiles across time. Sunbear can be used to impute single-cell temporal profile changes, align multi-dataset and multi-modal profiles across time, and extrapolate single-cell profiles in a missing modality. We applied Sunbear to reveal sex-biased transcription during mouse embryonic development and predict dynamic relationships between epigenetic priming and transcription for cells in which multi-modal profiles are unavailable. Sunbear thus enables the projection of single-cell time-series snapshots to multi-modal and multi-condition views of cellular trajectories.

## 1 Introduction

Rapidly improving single-cell sequencing technologies now allows us to characterize diverse and dynamic cellular processes. Multi-modal time series measurements of a population of cells can profile changes in gene expression, chromatin accessibility and structure, and DNA methylation as the cells carry out essential biological functions, differentiate, traverse the cell cycle, degenerate, or undergo stages of disease progression

Although such measurements have proven to be immensely valuable, they suffer from a fundamental limitation: because sequencing measurements are usually disruptive, it is impossible to trace a given cell’s behavior across time. In addition, single-cell profile inference suffers from heterogeneity of biological samples (e.g., female versus male, disease versus control) and data types (e.g., scRNA-seq, scATAC-seq), as well as limited sample availability. Consequently, a method capable of accurately imputing unobserved single-cell measurements—essentially answering questions such as, “What would the gene expression of this particular cell be if we had measured it at a different time point, or in a different experimental condition, or using a different measurement modality?”—would be immensely valuable in understanding the temporal dynamics and regulation of the behaviors of individual cells across biological conditions.

Many analytical techniques have been developed to model temporal processes from single-cell data. For example, pseudotime-based methods infer the ordering of cells along a one-dimensional or branching trajectory [46, 44, 38]. However, these methods do not perform imputation *per se*, and the relationship between the inferred pseudotime and actual time is left unspecified. Methods using RNA velocity to derive single-cell trajectories assume that nascent RNAs represent the future state of a cell [21, 23]. However, RNA velocity can be noisy and cannot be applied to other data modalities, and these methods are focused on ranking cells instead of predicting cellular profiles in missing time points.

In this work, we focus on methods that are specifically designed to integrate and extrapolate single-cell time series measurements (Table 1). For example, optimal transport-based methods, such as Waddington-OT [39], TrajectoryNet [45] and TIGON [40], operate on pairs of measurements in a time series, with the goal of aligning cells from two neighboring time points under the assumption that a single cell’s profile changes minimally between time points. Alternatively, neural ordinary/stochastic differential equation (ODE/SDE) based methods, such as PRESCIENT [51], RNAForecaster [9], MIOFlow [15], FBSDE [53] and scNODE [52], assume that each cell develops autonomously, and the cell’s future state is determined based on the cell’s current expression profile. Autoencoder models either assume that time is an additive variable in the embedding space [26] or are coupled with optimal transport or ODE based methods to optimize non-linear cell projection and cross-time alignment [45, 15, 52]. Graph-based methods, such as GraphFP [17], perform dynamic inference on top of cell graphs. However, most of these existing methods are focused on a single data modality (e.g., single-cell RNA-seq) or biological condition. Recently, Moscot.time was proposed to jointly perform optimal transport on multiomics data [18]. However, to our knowledge and as shown in Table 1, no existing method is designed to solve all of these problems at once.

**Table 1:** Methods for integrating and extrapolating single-cell time series data. OT = optimal transport; ODE = ordinary differential equations; AE = autoencoder.

|  | OT | ODE | AE | OT/ODE+AE | graph | multimodal OT | Sunbear |
| --- | --- | --- | --- | --- | --- | --- | --- |
| Optimized in the original data space |  |  | ✓ | ✓ |  |  | ✓ |
| Learns from distal time points |  | ✓ | ✓ | ✓ | ✓ | ✓ | ✓ |
| Does not require cell type labels | ✓ | ✓ |  | ✓ | ✓ | ✓ | ✓ |
| Corrects for batch effects |  |  |  |  |  |  | ✓ |
| Models multi-modal data |  |  |  |  |  | ✓ | ✓ |
| Performs imputation on missing conditions |  | ✓ | ✓ |  |  |  | ✓ |
| Performs imputation on missing modalities |  |  |  |  |  |  | ✓ |
| Citations | [39] | [51] | [26] | [45, 40, 9, 15, 53, 52] | [17] | [18] |  |

We thus set out to create a method that is capable of modeling various types of cellular profiles— gene expression, chromatin accessibility, DNA methylation, etc.—while capturing trends over time, between modalities, and across experimental conditions at single-cell resolution. We were particularly interested in the cross-modal setting where, for example, one data modality is “under-characterized”; i.e., we are missing measurements of that modality in one or more conditions or time points of interest. Although several existing methods have been developed for this type of cross-modal inference, they were developed for bulk analysis and have not been evaluated in the single-cell setting. For example, Sagittarius performs cross-species and cross-cell line inference of bulk transcriptomic profiles through a transformer model [48], and chronODE integrates bulk multi-omics time-series data to model the temporal dynamics of gene and chromatin features [5].

Following the lead of several existing methods [41, 25, 54], we hypothesized that a type of deep neural network known as a conditional variational autoencoder (cVAE) should be capable of accurately modeling a time series dataset consisting of multiple single-cell data conditions and modalities. Thus, our aim was to produce a model that would take as input a time series of single-cell measurements from two or more different data modalities and predict, for any cell in the input, its multi-modal measurement profile if the cell had been measured at some other time point (Figure 1A). To accomplish this goal, we created Sunbear, which is a cVAE that includes a continuous-time embedding and incorporates features within the model to enable cross-modality dataset matching (Figure 1B). To provide additional flexibility, Sunbear also includes latent embeddings to represent experimental or biological conditions (e.g., sex, batch, or perturbations). Intuitively, the model learns cell embeddings that represent cell identities while ensuring that these embeddings are conditionally independent of the time embeddings, study conditions, and batch effects. Critically, Sunbear is capable of learning non-linear relationships between time and cell identity, because the model is optimized to reconstruct accurate cellular profiles for all cells across all time points. Once the model is trained, we can vary the time factor continuously to infer smooth trajectories in cellular profiles across time. Furthermore, because Sunbear can be jointly optimized on multi-modal profiles, we can couple encoders of, for example, gene expression (scRNA-seq) with decoders of chromatin accessibility (scATAC-seq) to enable the prediction of missing scATAC-seq profiles across time (Figure 1C).

**Figure 1.**
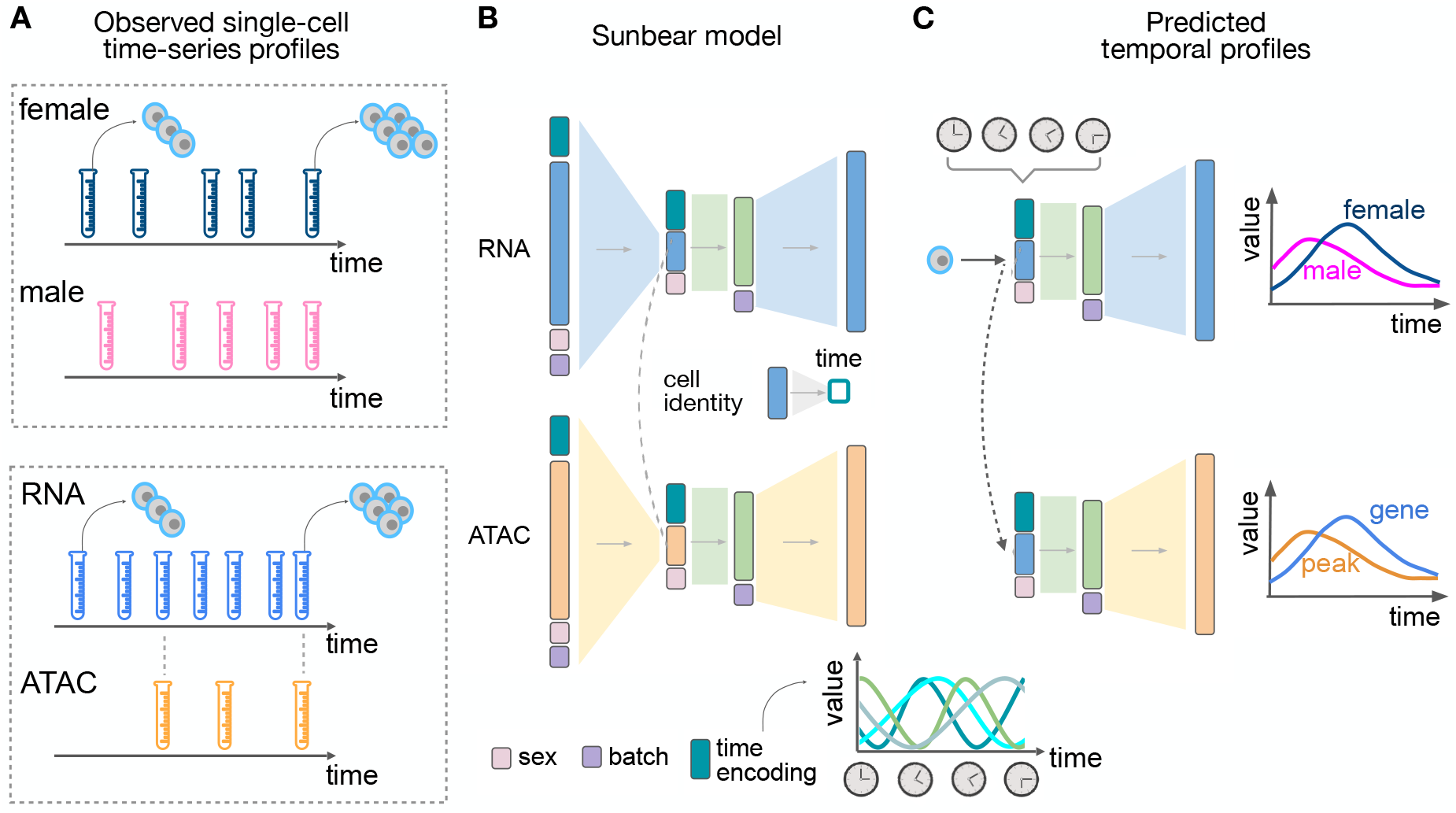
Sunbear Framework. (A) Sunbear takes as input a collection of measurements of single cells at multiple time points in two or more biological conditions (top) or data modalities (bottom). (B) SDuring the training phase, Sunbear learns to decompose the original time-series profiles into four components: cell identity, time point, batch and condition. Batch and condition factors are represented by one-hot encodings. The time factor is represented by a sinusoidal encoding. The cell identity factor is learned from the original profile and is conditionally independent of the other factors. In the multimodal setting, cell identities are aligned between data modalities. (C) In the prediction phase, Sunbear concatenates the query cell’s identity factor while varying other factors to impute the cell’s profile across time and condition. By transferring cell identity factors across modalities, Sunbear allows joint temporal modeling of multimodal profiles.

We apply Sunbear separately to time series scRNA-seq and scATAC-seq data collected during mouse embryonic development [33, 31, 1]. We first validate the accurate imputation capabilities of our model by predicting cellular profiles in held-out time points and conditions during embryonic developmental day E8 to E18.5. Next, we demonstrate that Sunbear enables identification of sex-biased transcription during mouse embryonic development and suggests a novel set of genes showing sex-biased transcription in nervous system and immune cells. Finally, we apply Sunbear to jointly model scRNA-seq and scATAC-seq data over mouse embryonic developmental days E7.5 to E8.75, showing that the model captures meaningful, time-relevant associations between transcriptional changes and chromatin priming during the formation of the hindbrain.

## 2. Results

### 2.1 Sunbear can predict a cell’s transcription profile at missing time points and missing conditions

We first set out to test Sunbear’s ability to predict a cell’s gene expression profile at missing time points. For this analysis, we used a set of one million cells randomly sampled from our recently published time series mouse embryonic development dataset, which consists of twelve million cells collected at 67 time points from embryonic day 8 through E18.75 (Figure 2A) [33]. To validate Sunbear’s performance at cross-time and cross-condition prediction, we held out one time point at a time and trained a model on the remaining time points (Section 4.5). In Sunbear, the time point information is encoded in a latent “time” factor, which is concatenated with the cell identity factor to impute the scRNA-seq profile of each cell. The model’s training procedure is designed to encourage the cell identity factors to be invariant of time. Therefore, by fixing the cell factors and varying the time factors, we can predict scRNA-seq profiles in a missing time point.

**Figure 2.**
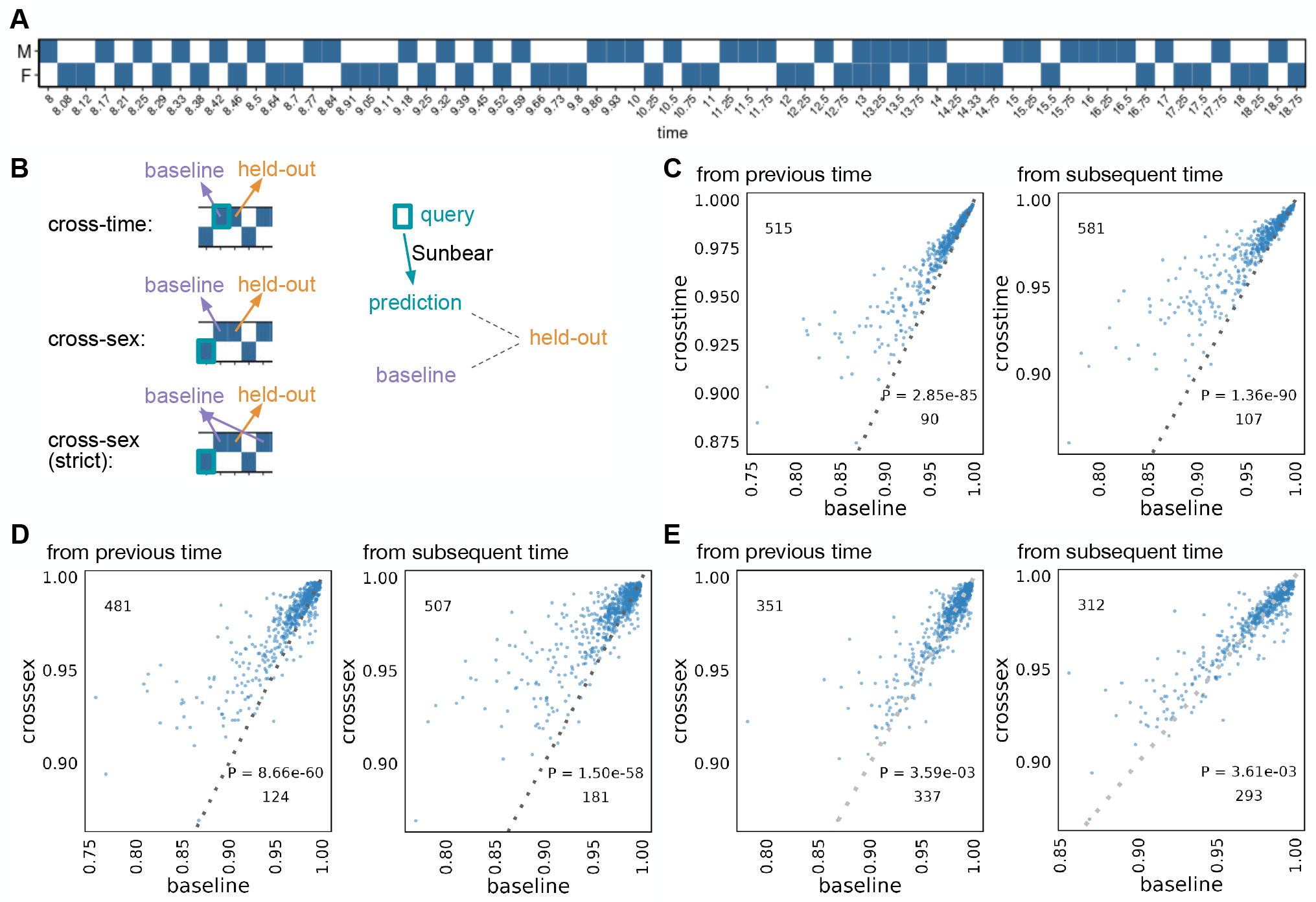
Single-cell profile inference across time and conditions. (A) Sunbear is trained on scRNA-seq profiles of whole mouse embryos collected at alternating sexes along developmental time points [33]. (B) Sunbear is validated in three scenarios. In each scenario, one data block is held out from the training, and Sunbear is used to predict the profile of the missing block based on cells in the query block (outlined in turquoise). Sunbear’s prediction is compared against the baselines using the held-out block’s nearest existing measurements with the desired sex factors. Pseudobulk Pearson correlation per major cell trajectory is calculated between the held-out profile and predictions/baselines. (C) Cross-time evaluation: query and baseline are selected either from the closest previous time point (left) or the closest subsequent time point (right). Pseudobulk Pearson correlation between the original held-out profile and predicted (y-axis) and baselines (x-axis) are plotted for each major cell trajectory in each held-out time point. Each dot represents a cell trajectory per held-out time point, and numbers indicate the number of dots above and below the diagonal line. P-values are calculated by a one-sided Wilcoxon rank-sum test. (D) Cross-sex prediction: similar to (C), except query cells are selected from the opposite sex to the held-out data. (E) Cross-sex prediction: similar to (D), except we enforce a strict baseline model by taking the mean of the previous and subsequent time point per cell trajectory.

To quantitatively evaluate Sunbear’s prediction performance, we would ideally like to measure Sunbear’s ability to predict how a single cell’s expression profile changes across time. However, because single cells are destroyed after sequencing, this type of evaluation is impossible. We therefore adopted an alternative approach that relies upon the cell type annotations created by the authors of the original study. These annotations allow us to ask how well each cell type-specific pseudobulk profile predicted in the missing time point agrees with the true cell type-specific pseudobulk profile that is held out from training (Section 4.5).

Specifically, to validate Sunbear’s performance on temporal inference, we asked whether the profile of each cell in the held-out time point can be recapitulated by using cells from the corresponding cell type at a neighboring time point. To do that, we compared Sunbear’s prediction (i.e., by varying the time factor of the query cell type at a neighboring time point to the held-out time point) with the baseline prediction, which is the pseudobulk profile of cells in the time point immediately before or after the held-out time point. Even though Sunbear did not see the cell-type information during training, it was able to predict scRNA-seq profiles in an unseen time point better than the baseline methods in the majority of cell types and time points (“cross-time”, Figure 2C).

In addition to modeling a continuous variable such as time, Sunbear is capable of modeling discrete factors corresponding to various types of experimental conditions. This functionality allows us to swap the condition factor to predict a cell’s profile in a missing time point and in a different condition, thus making direct comparisons of gene expression profiles across sexes. We used this functionality to predict sex-specific single-cell profiles at missing time points (“cross-sex”, Figure 2B). To evaluate our model’s performance, we held out each time point and sex at a time, and we used profiles from the opposite sex to recapitulate the held-out profile. Sunbear is able to predict scRNA-seq profiles in an unseen time point in the opposite sex, with significant improvement compared to using baseline models of the previous and/or subsequent time points in the held-out sex (Figure 2D–E).

### 2.2 Sunbear reveals sexual dimorphism in gene expression during mouse embryonic development

Having established Sunbear’s ability to capture sex differences in gene expression, we used the model to investigate sex-specific transcriptional differences during mouse embryonic development. Because only one sex is profiled at most time points, it is impossible to directly perform differential expression analysis between sexes at individual time points. Therefore, for each cell, we swapped the sex factor and calculated differential gene expression between sexes (Section 4.6).

We performed two types of validation of the transcriptional sex-difference predictions, taking advantage of the three time points (E13, E13.25, E13.75) for which the original data includes both female and male samples. Specifically, we performed evaluation on each cell type that has more than 1000 cells profiled across the three time points. First, we compared our prediction against the differential expression pattern between sexes calculated based on the original data at each sex-matched time point. For each of the three time points, we used the area under the receiver operating characteristic curve (AUROC) to assess the extent to which Sunbear’s sex-difference prediction identifies the female- and male-biased genes derived from the original dataset. As a baseline, we used the differential expression pattern from the closest time point as the sex-difference prediction for the time point of interest (Section 4.7). Comparing the two, we found that Sunbear significantly outperforms the baseline method (Figure 3A; One-sided Wilcoxon paired rank sum tests *p* = 1.24 × 10^−5^). In the second validation, we compared both the predicted and original sex difference patterns against a small set of genes known to escape X chromosome inactivation (XCI) in most/all mouse tissues (known constitutive escape genes; Section 4.7) [3]. Escape genes are transcribed from both copies of the X chromosome in females (XX), but only from the one X copy in males (XY), thus they are expected to have higher expression in females than males. Sunbear is able to rank the known escape genes to be more highly female-biased than other genes on the X chromosome (Figure 3B). In contrast, these genes are not ranked highly when we perform differential expression analysis directly on the original data (Figure 3B; One-sided Wilcoxon paired rank sum tests *p* = 8.3 ×10^−11^). These results suggest that Sunbear is able to capture time-specific and sex-biased transcription patterns.

**Figure 3.**
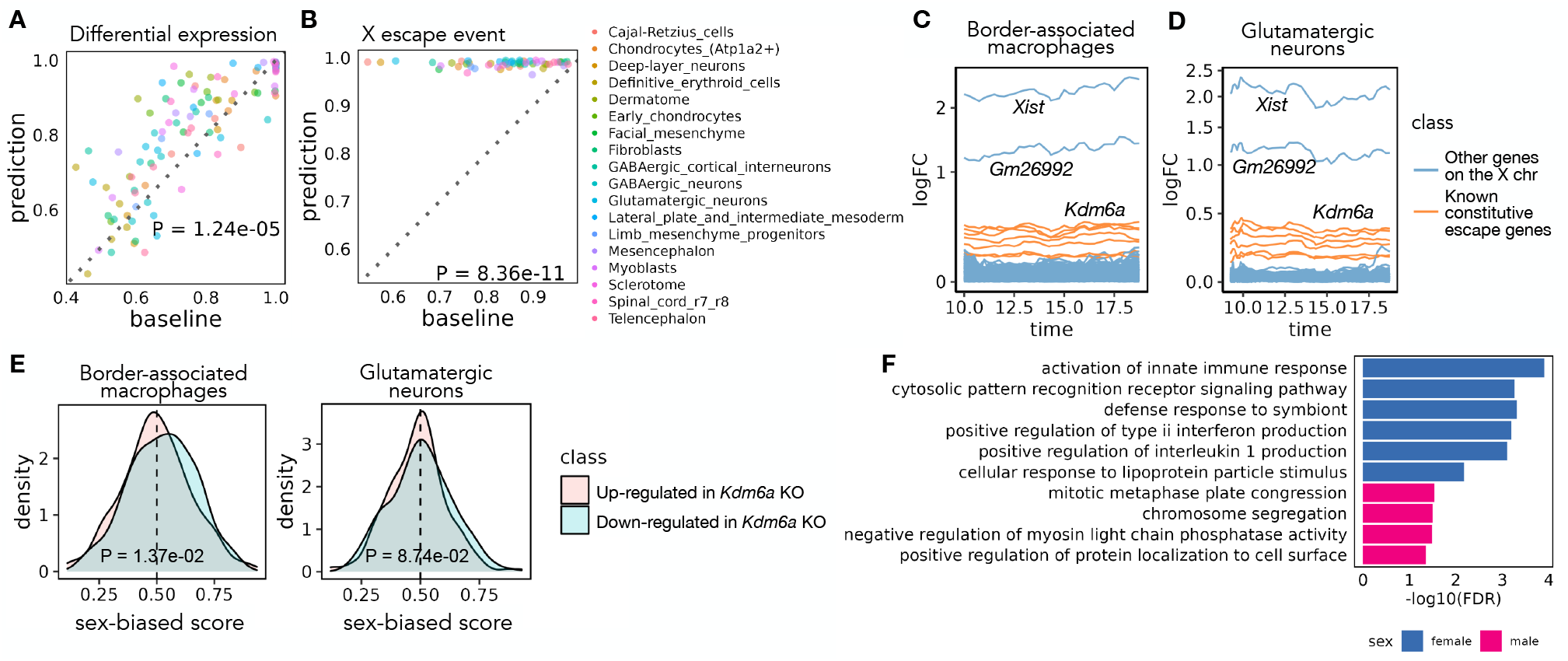
Sex differences in mouse embryonic development. (A) Pairwise comparison of Sunbear prediction and the nearest neighbor baseline in recapitulating differential expression patterns in sex-matched time points. Each dot indicates the AUROC score of recapitulating female/male-biased patterns in each sex-matched time point and cell type. (B) Similar to A, pairwise comparison of Sunbear prediction and the nearest neighbor baseline in ranking escape genes to be more female-biased than all other genes on the X chromosome. (C,D) Predicted temporal sex-biased log fold change in glutamatergic neurons and border-associated macrophages. Each line represents a gene that is predicted to be consistently higher in females than males and is colored by whether the gene is a known constitutive escape gene or not. (E) Distribution of predicted sex-biased scores of genes (0 = extremely male-biased, 1 = extremely female-biased), grouped and colored by whether the gene is up-(pink) or down-(blue) regulated in *Kdm6a* KO vs. WT samples in CD4+ cells. P-values are calculated by one-sided Wilcoxon rank sum tests. (F) Gene Ontology biological processes enriched in consistently female and male-biased genes in border-associated macrophages. Non-redundant terms with the smallest FDR are selected for visualization. No enrichment of biological processes is found in glutamatergic neurons.

We then systematically investigated the sex-biased pattern of two major cell trajectories across all time points: macrophages and central nervous system (CNS) neurons (Section 4.7). Specifically, we focused on the most abundant cell types within each trajectory: glutamatergic neurons and border-associated macrophages. Macrophages have been widely studied in the adult stage and exhibit well-known sex-biased transcription and phenotypes [12]. On the other hand, sex differences in CNS neurons could be limited since the brain is the somatic tissue showing the least sexual dimorphism in gene expression [50, 30, 37]. To our knowledge, no previous study has looked into sex differences in the expression of these cell types in vivo during embryonic development. Ranking Sunbear’s prediction based on the median log fold change between females and males across all time points, the top female-biased gene is consistently *Xist*, which is a gene expressed exclusively from the inactive X chromosome and is essential for the onset of XCI in females [24] (Figure 3C–D)). The second most female-biased gene is *Gm26992*, an antisense to *Xist* with an unknown function. This is followed by other constitutive escape genes, including *Kdm6a, Jpx, Eif2s3x, Pbdc1, Kdm5c, Ftx, and Ddx3x*. In addition to the 8-9 constitutive escape genes found in both cell types, macrophages show an additional 14 X-linked genes (out of a total of 356 genes with a female bias), while neurons only show 3 X-linked genes (out of a total of 226 genes with female bias) (Supplementary Table 1). This is consistent with the incomplete silencing of the X chromosome in macrophages [43].

Beyond sex-linked genes, Sunbear also predicts autosomal genes with consistent biases towards females or males in each cell type across time, although the predicted sex differences associated with these genes are associated with relatively small fold changes in gene expression (Supplementary Table 1), Figure 3C– D). Examples of autosomal genes with a female bias in macrophages include *Fcgr2b, Samd9l, and Ccr5*, which have been implicated in autoimmune diseases [47, 55, 35]. Fewer autosomal genes have consistent female bias in glutamatergic neurons, which show a few female-biased genes in common with macrophages (e.g. *Trim30a, Vmn2r29*). Interestingly, these genes show sex-biased transcription in embryos before sex hormones are produced, suggesting that they are regulated by sex-linked regulators within each cell.

We thus hypothesized that predicted female-biased autosomal genes are functionally associated with female-biased X-linked genes. To test this hypothesis, we focused on *Kdm6a*, which encodes a histone demethylase that removes the repressive histone modification H3K27me3 to activate gene expression and is predicted by Sunbear to be the escape gene with the largest female-biased expression among expressed escape genes (except for *Xist*) in multiple cell types. Specifically, we hypothesized that autosomal genes reported to be downregulated in a *Kdm6a* conditional knockout (KO) experiment in CD4+ cells (i.e., genes potentially activated by KDM6A) [16] would also show female-biased expression in border-associated macrophages compared to those upregulated in *Kdm6a* KO (i.e., genes potentially repressed by KDM6A). Indeed, Sunbear identifies the expected trend in border-associated macrophages (one-sided Wilcoxon rank sum test *p* = 1.37×10^−2^; Figure 3E). Meanwhile, we do not observe significant difference in neurons (*p* = 8.74×10^−2^), possibly due to low or no expression of these genes in neurons, or to different causes of sex bias in the two cell types.

Furthermore, we quantified the odds ratio of protein-protein interaction (PPI) between the consistently female-biased X-linked genes and autosomal genes using the STRING database (Section 4.8) [29], and observed a significantly larger number of PPIs between female-biased autosomal genes and X-linked genes in macrophages compared to the number of PPIs between random sets of autosomal genes and female-biased X-linked genes (permutation test *p* = 9.90×10^−3^; glutamatergic neurons are not included in this analysis because there are very few PPIs between sex-biased genes). These findings indicate that sex-biased transcriptional changes are not limited to sex-linked genes, and that autosomal sex-biased gene transcription may be influenced by sex-biased transcription of sex-linked genes.

Interestingly, although predicted consistent female- and male-biased genes in glutamatergic neurons are not enriched for specific biological processes, female-biased genes predicted in border-associated macrophages show significant enrichment in immune-related processes (hypergeometric test with FDR≤0.05, Supplementary Table 1; selected GO terms are shown in Figure 3F). These findings agree with previous evidence suggesting that immune processes tend to be stronger in females [19].

### 2.3 Cross-modality temporal profile inference reveals dynamic chromatin priming patterns

Sunbear can also be applied to infer multi-modal temporal profile changes. To do so, we trained Sunbear on scATAC-seq and scRNA-seq time-series profiles collected from overlapping time windows, including scRNA-seq (collected from E6.5-E8.5 with 6-hour intervals) [31] and scRNA-seq and scATAC-seq co-assays collected from four time points (i.e., E7.5, E8, E8.5, E8.75) [1]. We did not include the Qiu *et al*. dataset in this analysis because the authors staged embryos by somite count rather than embryonic developmental time between E8 and E10. Because scATAC-seq measurements are, in general, sparser and more expensive to generate than scRNA-seq measurements, we configured Sunbear to use scRNA-seq time-series data as a reference and trained with a similar strategy to the single-modality temporal inference model. The training of scATAC-seq model is guided by the scRNA-seq reference by encouraging colocalization of cell embeddings between corresponding cells in scRNA-seq and scATAC-seq, as well as by copying the time-relevant neural network layer learned from scRNA-seq to scATAC-seq. By doing so, Sunbear can jointly model time-relevant changes across data modalities (Figure 1).

We first asked whether Sunbear can recapitulate the temporal patterns observed in the under-characterized data domain (i.e., scATAC-seq) at a missing time point. To do that, we held out all scATAC-seq profiles from each embryonic developmental, and trained Sunbear on the remaining cells. As a sanity check, we first used UMAP to visualize the cell embeddings from different datasets, modalities and time points (Figure 4A, with E8 held-out), verifying that cells from different batches and data modalities are co-localized. Then, using scRNA-seq or scATAC-seq profiles in neighboring time points as queries (e.g., E7.5 or E8.5 for E8), we applied Sunbear to predict each cell’s chromatin accessibility in the held-out time point. Validating our prediction on the differential accessibility pattern between the held-out and each neighboring query time point per cell type, we found that Sunbear can accurately predict the temporal direction of chromatin accessibility changes (Section 4.9, Figure 4B). Interestingly, temporal accesbility can be recapitulated even when we use scRNA-seq profiles as queries. These observations demonstrate Sunbear’s ability to capture cross-modality temporal patterns in missing time points.

**Figure 4.**
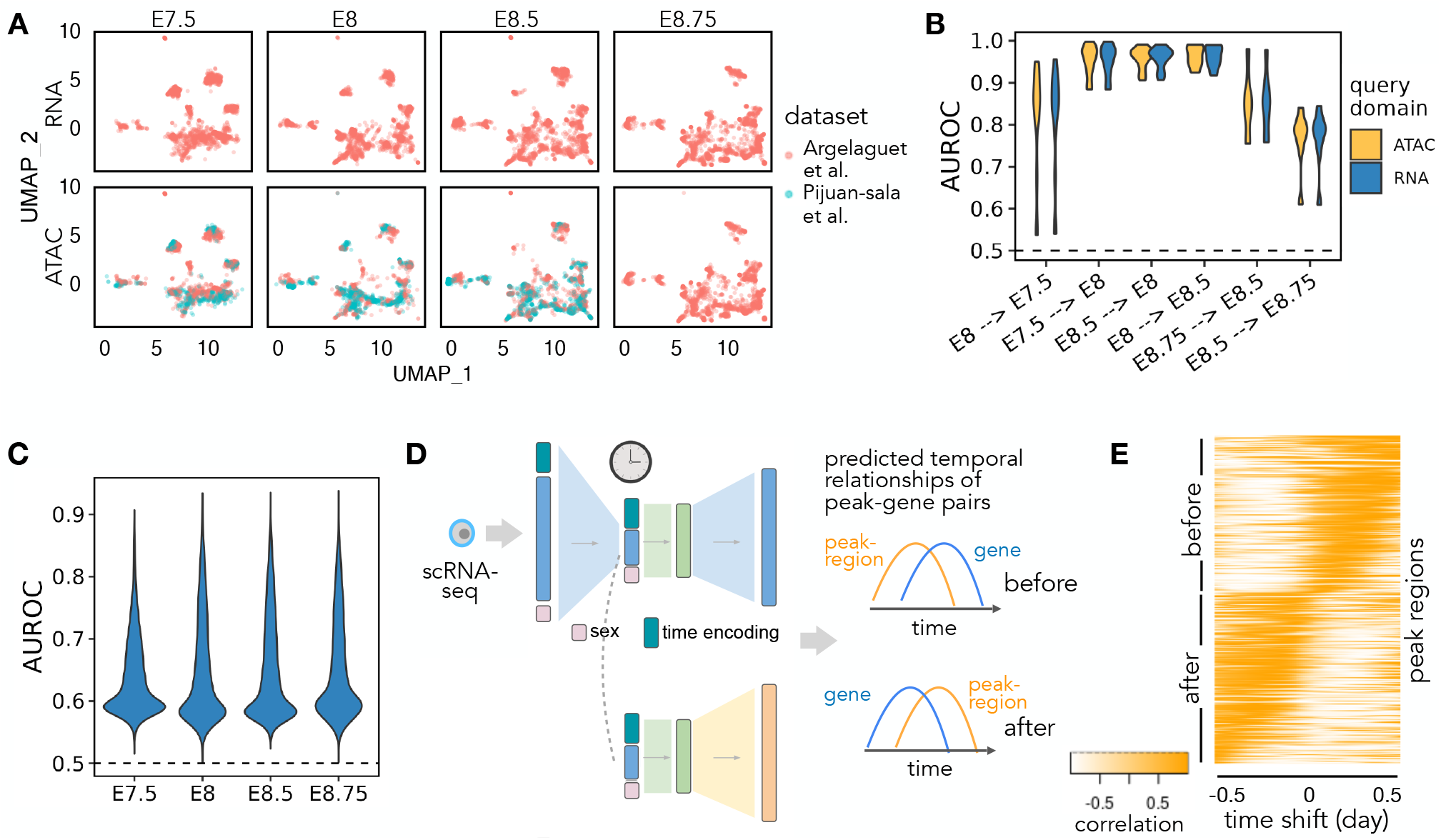
Multi-modal temporal inference. (A) A UMAP embedding suggests that scRNA-seq and scATAC-seq profiles are well aligned across time and batch. Only time points with both scRNA-seq and scATAC-seq available are shown. (B) AUROC of the predicted differential accessibility pattern relative to those derived from the original datasets. AUROC is calculated per cell type, and differential accessibility is calculated between each held-out time point and each query time point (shown as “query time point →held-out time point”). (C) Peak-wise AUROC of scATAC-seq profiles predicted based on scRNA-seq relative to the original scATAC-seq profile in each held-out time point. AUROCs are calculated across all cells. (D) Workflow for calculating the dynamic association between peaks and genes. A query cell’s scRNA-seq profile is fed into Sunbear to predict temporal patterns of gene expression and chromatin accessibility. For each pair of chromatin region and its proximal gene, we calculate the correlation coefficient between them with incremental time shifts, which results in a TLCC vector (column). (E) Predicted peak region-gene relationships. Heatmap of TLCC matrices on randomly selected 5000 peak regions with accessibility changes ahead (“before”) or subsequent to (“after”) nearby gene expression. Peak regions are sorted based on the time shift with the maximum TLCC.

Next, we assessed Sunbear’s ability to recapitulate cell-to-cell variations in the missing data modality (i.e., scATAC-seq) and time point. To do that, we used scRNA-seq profiles from the held-out time point as queries to predict each cell’s corresponding chromatin accessibility profiles. We then compared the predicted profile against original scATAC-seq measurements by comparing the predicted and true scATAC-seq profiles per peak region across all cells (Figure 4C), and found accurate prediction of accessibility trends in more than 99.9% of peaks across all time points (Section 4.9).

Having validated Sunbear’s performance on multi-modal temporal inference, we trained the model on all time points, with the aim of characterizing patterns of temporal coordination between transcription and chromatin accessibility changes within each cell. Here, we focused on the hindbrain trajectory around E8.25, when the cell lineage first emerges [33]. Because there are no scATAC-seq profiles available at E8.25, we used scRNA-seq profiles collected in E8.25 in the hindbrain trajectory as queries to predict temporal cellular transcription and accessibility profiles (Figure 4D). We then used time lagged cross correlation (TLCC) analysis [6] to quantify the dynamic coordination between the predicted gene transcription and its nearby region’s accessibility changes (Figure 4D). Sorting genomic regions based on their temporal association pattern with nearby genes, we found diverse dynamic coordination between gene expression and chromatin priming (Figure 4D–E, Section 4.10). In particular, we observe a continuous gradient of time lags between chromatin accessibilities and their nearby gene expression changes. These patterns can be categorized as peak regions changing before or after gene expression changes.

We further asked whether distinct sequence motifs are enriched in genomic regions where Sunbear predicts peak accessibilities change before and after transcription changes. This analysis showed that several transcription factor binding motifs are enriched in genome regions in the “before” category (e.g., CTCF, Zic2, and Zic3), but none in the “after” category (details in Section 4.10). Given that transcription initiation requires the opening of chromatin regions at enhancer and promoter regions and the recruitment of transcription factors, our finding agrees with a model in which chromosomal regions tend to be primed with transcription factors before transcription changes [4, 22, 34]. Among the transcription factors that we predicted to prime chromatin regions before transcriptional changes, Zic2 and Zic3 are important transcriptional regulators of hindbrain segmentation [8, 7]. Meanwhile, CTCF is a well-known mediator for chromatin loops and has been suggested to indirectly affect the regulation of genes involved in hindbrain development [11]. Overall, the above evidence suggests that Sunbear is able to identify temporal coordination between chromatin priming and transcription based on static snapshots of cells measured in one modality, and that Sunbear can offer insights into transcription factors involved in lineage priming during development.

## 3 Discussion

A critical question for characterizing single cells is to understand how each cell’s profile changes across time in vivo. While most methods focus on inferring cell trajectories across time points collected from a single modality or condition, questions remain about how to model continuous profiles, compare profiles across biological conditions with unmatched time points, and understand the coordination between multi-omic features. Sunbear provides a deep learning framework for temporal profile inference across data modalities and conditions. The Sunbear model jointly integrates datasets collected from different time points, batches, conditions and modalities, and the model outputs continuous cellular profile changes across time. Sunbear thus allows us to directly compare cellular profiles across conditions and investigate joint temporal relationships between modalities. Furthermore, Sunbear can be trained across millions of cells and the full space of measured features (e.g., genes and peaks).

Studies of sex differences during embryonic development are few and often lack matched biological samples across time points. In addition, differential expression analysis at the cell type level may suffer from biases due to differences in sub-cell type composition between conditions. Our method mitigates this bias by computationally inferring cellular profiles at missing time points and conditions and by performing cross-condition comparisons at single-cell resolution. Our predictions, for the first time, systematically characterized sex-biased expression patterns during mouse embryonic development. Besides recapitulating well-known sex-biased genes, we generated various data-driven hypotheses for downstream experimental validation in the future.

Sunbear also identifies dynamic associations between chromatin priming and transcription. Previous studies on multimodal measurements have found or assumed that chromatin priming tends to occur ahead of transcription changes [27, 49]. In our analysis, we instead focused on the scenario where only one type of data modality is measured, and we computationally predicted the multimodal dynamic coordination in each cell. Our results not only agree with previous knowledge but also suggest diverse types of dynamic coordination (e.g., different time delays, as well as positive and negative correlations). With the increasing amount of effort being devoted to characterizing mouse embryonic development, we envision that our temporal predictions can be combined with genome annotations from corresponding cell types and time points in the future.

In the future, Sunbear could be improved in several ways. First, although our model does not fully rely on the assumption of consistent cell proliferation or death rate, incorporating an explicit model of cell death and cell proliferation could further improve the temporal alignment, as was done in [51, 39]. Second, as a proof of concept analysis, our current multi-modal model leverages some co-assay data to guide the alignment of cells across modalities. Generalizing the framework to carry out cross-modality alignment without requiring co-assay data would greatly enhance the applicability of Sunbear.

In the future, we envision the Sunbear framework being applied to integrate and compare human disease conditions collected at varying time points, characterize sequential transcription factor activation, and reveal the dynamic multi-omic coordination during cell fate specification.

## 4 Methods

### 4.1 Mouse embryonic development data

The single-cell RNA-seq datasets used in this paper are listed in Table 2. For each dataset, we obtained the raw count-by-cell matrices. For the co-assay dataset from Argelaguet et al., only time points with at least two biological replicates in this dataset were retained. For efficiency of model training, we randomly subsampled cells in the Qiu et al. dataset to retain one million cells. Because P0 samples exhibit a huge environmental shift relative to embryonic stages, the P0 samples are not included in this analysis. In addition, because embryos before E10 are staged by somite counts, we converted the somite number to developmental time points by assuming equal time intervals between somite stages (i.e., each somite stage has incremental time of 2(day)/34(somite)).

**Table 2:** Time-series data in mouse embryonic development.

| Assay | Strain | Sex | Start | End | Freq | Source |
| --- | --- | --- | --- | --- | --- | --- |
| sci-RNA-seq | BL6 | m&f | E8 | E18.75 | 2-6h | Qiu et al. [33] |
| sci-RNA-seq | BL6 | - | E6.5 | E8.5 | 6h | Pijuan-sala et al. [31] |
| co-assay: scRNA-seq & scATAC-seq | BL6 | - | E7.5 | 8.75 | ~6h | Argelaguet et al. [1] |

### 4.2 Modeling scRNA-seq temporal profiles

The goal of Sunbear is to decompose observed cellular profiles into latent factors representing cell identity, batch or condition, and the time point when each cell is collected. In this way, we can predict a cell’s profile across time and conditions by concatenating its cell identity factor with varying time and condition factors. Thus, Sunbear takes as input single-cell time-series profiles and reconstructs them by cell identity, time and batch/condition factors. Specifically, to interpolate continuous shifts of single-cell profiles across time points that are not captured, Sunbear represents time with sinusoidal encodings. I.e., the embedding for each time point *t* is a *d*-dimensional vector, represented as

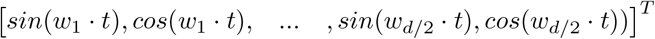

where *k ∈*0, 1, …, *d ™*1 and *w*_*k*_ = 2*π/*10000^2*k/d*^, *t* is represented in the units of day.

Sunbear is optimized toward two goals: obtaining accurate reconstruction of the observed single-cell profiles and aligning cells collected across different time points. To achieve these two goals, Sunbear adapts the conditional variational autoencoder (cVAE) framework [25, 54], with conditions as sinusoidal-encoded time factor and one-hot encoded sample conditions (e.g., sex, batch). Sunbear assumes that scRNA-seq counts follow a zero-inflated negative binomial (ZINB) distribution and estimates the reconstruction loss as the log-likelihood of the ZINB distribution [25]. Given scRNA-seq from time points 1 to *T*, Sunbear minimizes the following loss function:

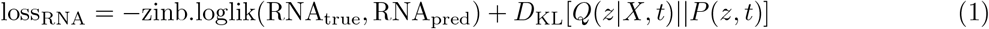

To further encourage cell embeddings to be independent of time, we also add a discriminator that tries to predict time points based on cell embeddings, and we perform an alternative training by iteratively minimizing the discriminator loss:

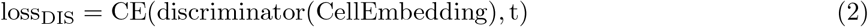

and the generator loss:

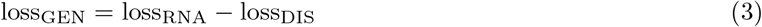

During model training, we held out all cells from one time point as the test set. Validation and training sets were created by randomly splitting the cells in the remaining time points with a 1:4 ratio. When there were large numbers of cells or time points, to make sure the model does not overfit to a high abundance time point, we capped the number of cells in the validation set at each time point to be 2000. Once Sunbear is trained with the existing time points, we can estimate single-cell profiles for any missing time point by swapping the time vector to the corresponding time point. We can also swap the condition factor to predict a cell’s profile if it were captured in another data condition (e.g., an opposite sex).

### 4.3 Extension of Sunbear for multi-modal temporal infererence

Next, we extended Sunbear to jointly infer temporal profiles across data modalities based on single- and multi-modal time-series profiles. Similar to the single-modality case, for each data modality, Sunbear learns cell embeddings that represent cell identities, which are conditionally independent of the time embeddings, conditions and batch effects. For the scATAC-seq domain, we binarize the scATAC-seq profiles and assume that they follow a Bernoulli distribution, estimating the reconstruction loss using binary cross entropy [2]. Thus, the scATAC-seq model is optimized with

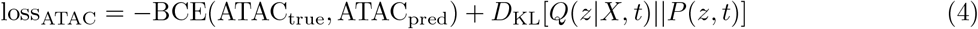

Because of the imbalanced time series measurements across data modalities (e.g., scRNA-seq time series usually cover more time points than scATAC-seq), we designed Sunbear to take one data modality with denser time points as a reference to guide the imputation of the data modality with sparser time points. To this end, we optimized Sunbear by using scRNA-seq as a reference and forced the scATAC-seq model to have a shared, time-relevant network layer and cell embeddings with the scRNA-seq model. By doing so, we enable the sharing of interaction effects among cells, time and conditions across data modalities.

Sunbear does stepwise optimization during training. In the first step, we train the scRNA-seq time model with Equation 1, while fixing all weights in the scATAC-seq model. In the second step, we train the scATAC-seq time model with regard to the scRNA-seq reference (Equation 5), with weights in the time factors fixed to be the same as the scRNA-seq, and with the scATAC-seq cell embeddings forced to fall close to the scRNA-seq cell embeddings. We leverage the one-to-one cell correspondence in co-assay datasets to supervise the alignment of cell embeddings between data modalities. This is achieved with a mean-squared error (MSE) loss between corresponding cell embeddings, together with a translation loss between scRNA-seq and scATAC-seq co-assay profiles:

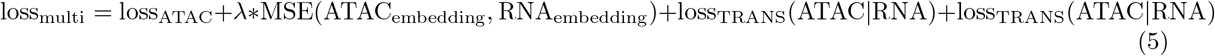

During training, Sunbear iteratively optimizes the scRNA-seq model and the scATAC-seq model, so that the scATAC-seq model can slowly adapt to the cell embeddings and shared interaction effects of the reference scRNA-seq model.

To validate the model, we held out all single cells from one time point as the test set, and we randomly split cells from the remaining time points into a validation set (1/5) and a training set (4/5). The number of cells in the validation set at each time point is capped to be 2000.

### 4.4 Hyperparameter tuning

As with any deep learning model, Sunbear has a variety of hyperparameters that need to be tuned. Accordingly, we performed a grid search on the following hyperparameters:

- number of latent dimensions in the VAE ∈ {25, 50, 100}
- minimum wavelength of sinusoidal encoding ∈ {1, 2π}
- MSE weight *λ* (when trained on multi-modality data) ∈ {1, 100, 10000}

The number of hidden layers was fixed at 2, and the dimension of the time embeddings was fixed at 50. We trained one model for each of the 3*×*2*×*3 = 18 points in our hyperparameter grid, and we selected the best-performing model based on the validation set. Ideally, the model should be able to reconstruct the observed profiles accurately using the neighboring time points, without relying on cell type labels. Thus, we calculate the cross-time prediction’s pseudobulk Pearson correlation with the original profile, as well as the LISI score [20] between neighboring time points, as ways to assess the ability of each model to generalize across time. In the single modality model, we select hyperparameters based on the summed rank of cross-time pseudobulk Pearson correlation, LISI score across all neighboring time points, as well as LISI score between neighbors of the held-out time point. When multiple batches are available, we also include the LISI score between batches in the model selection criteria. In the multi-modality model, we select the best hyperparameters using the above criteria based on the non-reference (i.e., scATAC-seq) data modality, together with the cross-modality LISI score and cross-modality translation losses in both directions.

### 4.5 scRNA-seq temporal inference evaluation

We evaluate the model’s ability to generalize to uncharacterized sex and time points through leave-one-out cross-validation; i.e., for each time point and each sex, we hold out the entire cell population from training and predict its profile from neighboring time points (“cross-time”) or the opposite sex (“cross-sex”). Because of the disruptive nature of single-cell measurements, there is no ground truth for a single cell’s profile at another time point. Therefore, for the purpose of validation, we instead leverage cell type labels, as previously annotated by biological experts, and ask whether the pseudobulk gene expression of each cell type can be correctly recapitulated, compared to pseudobulk profiles from corresponding cell types in neighboring time points.

To evaluate at the cell type level, for each cell type in the held-out time point, we predict its profile using the neighboring time point’s profile in the same cell trajectory and compare the prediction accuracy using pseudobulk Pearson correlation between the held-out observed value and the predicted value. Specifically,because our prediction corrects for the sequencing depth of single cells, to make a fair comparison, we calculate the pseudobulk profile by normalizing every single cell’s sequencing depth first before averaging them across cells. Only cell types with at least 25 cells in the held-out time point are considered in this evaluation.

For each held-out sex and time point, we make predictions with four combinations of starting points: previous or subsequent time points in the corresponding (“cross-time”) or opposite sex (“cross-sex”). The performance in each of these settings is compared against a corresponding baseline, which is the pseudobulk Pearson correlation of the held-out cell type with the cell type-specific profiles in the corresponding previous or subsequent time points in the corresponding sex. We also compared Sunbear to a stricter baseline: the pseudobulk profiles of the corresponding cell trajectory in the previous and subsequent cell types. This is a difficult baseline to beat because the cell trajectory is annotated with regard to the evidence that the previous, held-out and subsequent cells in the same cell type should follow a sequential trajectory. However, Sunbear has not seen cells in the held-out time point or any cell type labels, and is forced to make predictions only based on the previous or subsequent time point. Given that embryos collected before E10 are staged by somite number, where clock time cannot be precisely determined, we only perform such evaluation for samples collected after E10. One-sided Wilcoxon paired rank sum tests are conducted to compare the prediction performance with baselines.

### 4.6 Predicting transcriptional sex differences across time

For each gene, Sunbear returns a sex-biased score based on sex-matched predicted profiles across cells within each cell type. To do that, first, we predict each cell’s denoised profile in both sexes. Then, for each cell type, we calculate each gene’s sex-biased pattern over time by performing, for each time point, a one-sided Wilcoxon signed-rank test between predicted gene expression in females and males across all cells. Along with each statistical test, a sex-biased score is calculated as the sum of positive ranks divided by the sum of all absolute ranks *n*(*n* + 1)*/*2. We rescale the score to the range [0, 1], with values close to 0.5 indicating minimal sex difference and scores of 1 and 0 representing strongly female- or male-biased expression. To make sure the sex-biased is robust to outliers, we only calculated such a score when there are more than 50 cells within a cell type.

### 4.7 Validation of sex-difference prediction

To validate Sunbear’s sex-difference prediction, we first retrieved sex-biased transcriptional patterns from three time points (E13, E13.25, and E13.75) with matched female and male samples in [33]. Specifically, 18 cell types with enough observations across these three time points (i.e., *≥*1000 cells) are selected for downstream evaluation. Because there are no biological replicates available at each time point, following the suggestion in [42], for each cell type, we calculated the original differential expression pattern between sexes by applying a Student’s t-test on sequencing-depth corrected and log-normalized single-cell profiles. Genes expressed in less than 5% of cells are excluded from the differential expression calculation. Genes with Benjamini-Hochberg corrected FDR≤ 0.05 are labeled as significantly up- or down-regulated genes.

We then validated Sunbear’s sex-difference prediction against the original differential expression pattern within each time point and cell type. To mimic the scenario where only one sex is profiled at a time point, we held out the male sample in the time point of interest from model training, and then calculated sex-biased scores based on the predicted profiles. We tested female- and male-biased expression prediction separately. In each direction, genes that are significantly female-/male-biased (FDR≤ 0.05) are assigned positive labels and all other genes are labeled as negatives. AUROC is calculated for our prediction against differential expression labels. In the end, for each cell type and time point, two AUROC scores are calculated.

As a baseline, we calculated the AUROC of the original differential expression patterns in the closest sex-matched time point against the female/male-biased gene labels in the time point of study. A one-sided Wilcoxon paired rank sum test was conducted to compare the prediction performance between Sunbear and the baseline across all matched time points, cell types and differential expression directions.

Besides comparing Sunbear’s prediction with the original measurements, we also validated the predictions against prior knowledge of sex-biased transcription. To do so, we retrieved X-escape genes (including *Ddx3x, Kdm6a, Kdm5c, Eif2s3x, Pbdc1, Jpx, Ftx and 5530601H04Rik*) from [3] and calculated the AUROC of predicting these genes to be more female-biased than all other genes on the X chromosome. For this analysis, we used the differential expression pattern between sexes in original measurements as a baseline. For each cell type and time point, AUROCs based on our prediction are compared against those based on baselines using a one-sided Wilcoxon paired rank sum test.

### 4.8 Sex-biased gene and pathway analysis

To systematically characterize sex differences across time and avoid biases introduced by individual samples, we used two additional strategies to ensure robustness. First, we trained ten models with a random time point held out from training each time, and we took the median of sex-biased statistics across these models as the predicted sex-biased score. Second, instead of comparing sex differences based on one biological sample at each time point, we calculated sex differences across cells from the current, previous and subsequent time points. These approaches allow us to derive robust sex-biased calculations across multiple biological samples. To identify sex-biased genes for each cell type, we focused on time points with more than 25 cells in that cell type. In order to focus on genes that are important in that cell type, for each time point in that cell type, we ranked genes by predicted mean expression level across sexes and retrieved the first half of highly expressed genes. Only genes that are consistently expressed across all time points in the corresponding cell type are included in downstream analysis. Sex-biased genes are determined as genes with sex-biased scores consistently falling above or below 0.5 across all time points. GO term enrichment analysis was performed using a hypergeometric test, and only those terms with Benjamini-Hochberg corrected FDR≤ 0.05 are reported.

Protein-protein interaction information from *mus musculus* was downloaded from the STRING database [29]. Because we are interested in the broad definition of interactions (i.e., including coexpression, shared pathways, etc.), we did not apply additional filtering to the score and type of interaction in the STRING database. The odds ratio was calculated as the ratio of the odds of interactions happening between female-biased autosomal and X-linked genes, compared to odds of female-biased X-linked genes interacting with non-female-biased autosomal genes. Statistical tests are only carried out when there are *≥* 50 PPIs. We then generated the null distribution by calculating the odds ratio of interactions between randomly sampled autosomal genes and female-biased X-linked genes. Random sampling is repeated 100 times, and the size of the random sample is equal to the number of female-biased autosomal genes.

### 4.6 Multi-modal temporal inference evaluation

We first tested whether Sunbear can recapitulate temporal changes in scATAC-seq at a missing time point. Because we do not have ground truth data at the single-cell level, we instead asked whether Sunbear can correctly predict the direction of chromatin accessibility changes across time in each cell type. To do that, we held out all scATAC-seq profiles from one time point and used scRNA-seq or scATAC-seq profiles at a neighboring time point as queries to predict the missing time point’s scATAC-seq profiles. For each cell type and each peak, we predicted peak accessibility changes between the missing time point and the query time point using the test statistic of the one-sided Wilcoxon paired rank sum test (i.e., the sum of positive ranks divided by the sum of all absolute ranks). To make sure the prediction is robust to random initiation and data split during model training, we trained ten models with different random seeds and took the median of differential peak assessibility statistics as the final prediction. We then compared our prediction against differentially accessible peaks between cells measured in the held-out and the query time point within the corresponding cell type. Only cell types with *≥*25 cells in both time points were retained. Differential accessibility of the original profiles was calculated using edgeR on the pseudobulk level [36, 10]. Finally, we report the AUROC of our prediction against significantly up- and down-regulated peaks (labeled as positives and negatives, FDR ≤0.05).

To validate Sunbear’s performance at capturing individual-cell-level differences in the missing data modality, we used scRNA-seq profiles in the held-out time point as queries, and we asked how well their corresponding chromatin accessibility profiles can be recapitulated. For each genomic region that is accessible in more than 5% of cells in the held-out population, we calculate the AUROC of the predicted accessibility against the original binary accessibility pattern across all cells. Similarly, for each genomic region, we also calculate the p-value of its predicted accessibility to be higher in cells with peaks in the original measurement than those without, using a one-sided Wilcoxon rank-sum test. The fraction of peaks with Benjamini-Hochberg corrected FDR≤0.05 are reported.

### 4.10 Identifying temporal dynamics between chromatin priming and transcription

To investigate the dynamic relationship between gene expression and chromatin accessibility, we varied the time factor for a query cell to predict continuous changes in transcription and chromatin accessibility. We chose to focus on the hindbrain trajectory since it first emerges at E8.25, when scATAC-seq profiles are not available. In addition, the hindbrain trajectory is conserved across vertebrates, thus enabling us to validate our results on findings derived from model organisms (e.g., zebrafish) [13]. We also focused on genes with significantly increased or decreased expression levels in the hindbrain at E8.25 relative to its parent trajectory at E8. We reasoned that temporal regulation of these genes is likely critical for hindbrain formation [32]. For each gene, we retrieved its proximal peaks that are mapped to 200kb flanking regions upstream of its transcription start site. Then we predicted the gene expression and proximal chromosome accessibility changes within a [7.5, 9] day interval, and we calculated the time lagged cross correlation (TLCC) of the gene with respect to each of its proximal peaks regions.

The TLCC score is calculated via the following steps. For each peak region and gene pair, we shift the predicted accessibility profiles along the time axis at 0.01-day steps, considering shifts in the range [-0.5, 0.5]. Then, for each step, we calculate the Pearson correlation between the predicted expression profile and the shifted temporal accessibility profile within the core [8, 8.5] day interval. Thus, each peak region and gene pair in a cell can be represented as a 101-dimensional vector consisting of Pearson correlations corresponding to the varying time shifts. Concatenating all pairs of genes and peak regions, we obtain a TLCC matrix with 101 by #region-gene pair dimensions.

To maintain single-cell resolution while avoiding potential noise generated from a single cell or a single model, we trained a set of models with ten random seeds and took the median of the predicted gene expression values or peak accessibility values. For a query cell, we selected its four closest neighbor cells summarized across the ten models. Specifically, for each model, we retain its top 25 nearest neighbors based on Euclidean distance on Sunbear’s cell embeddings and then sum the ranks across ten models to obtain the final ranking. Cells that are not among the top 25 neighbors in any of the runs are excluded from the nearest neighbor list. On top of the five cells, we used Sunbear to predict each cell’s multimodal temporal TLCC matrices and concatenated the TLCC vectors across the five cells along the peak gene axis. Because not all proximal chromatin regions regulate transcription, we removed region-gene pairs with a maximum correlation of less than 0.5 and only retained peak region and gene pairs showing up in more than one cell after the filtering step. Finally, we categorize peak regions into two categories: those whose accessibility pattern consistently changes ahead of its nearby gene expression pattern (“before”) or subsequent to the gene expression pattern (“after”).

To identify sequence features specific for each category, we first retrieved all unique sequences from peak regions within each category and then used MEME-ChIP to generate enriched motifs for one set of genomics regions against the other (with “Differential Enrichment mode” mode) with default cutoffs [28]. Tomtom was used to identify transcription factors with q-value ≤ 0.05 [14].

## Data availability

All single-cell data used in the paper are downloaded from the public database. sci-RNA-seq data from E6.5 to E8.5 are downloaded from https://tome.gs.washington.edu/ [32]. sci-RNA-seq data from E8 to E18.75 are downloaded from omg.gs.washington.edu/ [33]. scRNA-seq and scATAC-seq co-assay data from E7.5 to E8.75 are retrieved from GSE205117 [1].

## Code availability

The Apache-licensed Icebear source code is available at https://github.com/Noble-Lab/ Sunbear.

## Author information

### Contributions

Ran Zhang: Conceptualization, Methodology, Formal analysis, Investigation, Validation, Visualization, Writing - Original draft preparation, Software. Chengxiang Qiu, Gala Filippova, Gang Li: Investigation, Validation. Jay Shendure: Supervision. Jean-Philippe Vert: Supervision, Methodology. Xinxian Deng, Christine Disteche: Supervision, Investigation, Validation. William Stafford Noble: Conceptualization, Supervision, Methodology, Funding acquisition. All authors paticipated in the Writing - Review & Editing process.

## Ethics declarations

The authors declare that they have no conflict of interest.

